# Frequency and Prognostic Value of NPM1 Mutations in Sudanese Acute Myeloid Leukemia Patients

**DOI:** 10.1101/2020.05.31.126334

**Authors:** Eman Ali Elzain, Hiba BadrEldin Khalil

## Abstract

**Introduction:** Acute myeloid leukemia (AML) is a malignancy of proliferative, clonal, abnormally, or poorly differentiated cells of the hematopoietic system, characterized by clonal evolution and genetic heterogeneity (Döhner *et al*, 2015). The molecular genetics of AML regarding the prognosis of patients mainly representing by NPM1. NPM1 is a nucleolar protein that located on chromosome **5q35.1** which shuttles between the nucleus and cytoplasm. It is concerned in multiple functions, including ribosomal protein assembly and transport, control of centrosome duplication, and regulation of the tumor suppressor ARF & p53. NPM1 mutations that relocalize NPM1 from the nucleus into the cytoplasm are associated with development of acute myeloid leukemia (Garzon *et al*, 2008). The aim of this study was to determine the frequency and clarify the prognostic value of NPM1 mutations among Sudanese patients with acute myeloid leukemia.

**Materials and Methods:** A cross sectional study recruited 100 patients in this study clinically diagnosed firstly (not transformed from any other malignancy) as AML patients based on the diagnostic protocol concern RICK hospital; such as morphological identification, immunophenotyping analysis, and molecular genetics, Of the 100 AML patients, 54% were newly diagnosed and 46% were admitted by chemotherapy treatment and follow up. EDTA blood or bone marrow samples were collected from patients for performing CBC, hematological studies including FAB classification, PCR protocols and sequencing. genomic DNA was extracted from all samples using guanidine method (Ding et al, 2008).

**Results:** NPM1 mutations detection, sequencing technique was done, and then sequencing analysis by software (Mega 7 Software) (Kumar *et al*, 2016) revealed that there were no NPM1mutations in Sudanese AML patients.

**Conclusion:** The Sudanese AML patients carry wild type NPM1.

## 1. Introduction

Acute myeloid leukemia (AML) is the most familiar hematologic malignancy, characterized by uncontrolled construction of hematopoietic stem cells follow-on in abnormal accumulation of myeloblasts. Generally, based on the cytogenetic abnormalities, the prognosis of AML patients is grouped into three risk groups: good, intermediate, and poor (Rezaei *et al*, 2017).

Acute myeloid leukemia is the result of more than one mutation; one of the most common is NPM1mutations. NPM1 mutations represent frequent genetic alterations in patients with AML associated with a favorable prognosis. Moreover, NPM1 mutations displays distinct biological and clinical features that led to its inclusion as a provisional disease entity in the 2016 World Health Organization (WHO) classification of myeloid neoplasms (Arber et al, 2016). Majority of cases NPM1 mutations result in a frameshift due to an insertion of four bases, which gather in exon 12. Different types of NPM1 mutations have been illustrated according to the inserted tetranucleotide, the most frequent being type A mutations (TCTG) in 80%, followed by type B (CATG) and type D (CCTG) mutations in about 10%, and a spectrum of other mutations accounting for 10% of cases. In rare cases, insertions from 2 to 9 bases can occur (e.g. types E, F). The altered NPM1 protein (NPM1c+) contains an additional C terminal nuclear export signal (NES) motif and loses at least one tryptophan residue, causing an aberrant cytoplasmatic localization of the protein (Pastore *et al*, 2014). its roles, it relies on its ability to shuttle between the nucleolus, nucleus and cytoplasm using subcellularlocalization signals. This ability is impaired in 30% of AMLs as a result of NPM1c mutations, which disrupt the nucleolar localization signal of NPM1 and generate a nuclear export signal in its place. Mutant NPM1 is known to bind to and alter the subcellular distribution of several proteins, including HEXIM1, p19Arf and nuclear factor-kB;8 however, the relevance of these interactions to AML is unclear (Mupo *et al*, 2013), The intrinsic genetic heterogeneity of AML, which has been well described, includes a sizeable minority whose blasts have NPM1 mutations (Papaemmanuil *et al*, 2016). AML patients whose blasts have an NPM1 mutation but do not have a FLT3 internal tandem duplication (ITD) represent a common molecular category and have a relatively good prognosis (Uckelmann *et al*, 2018). So this study was designed to determine the frequency and prognostic value of NPM1 mutations among Sudanese AML patients, Moreover, according to our knowledge there was no published data about NPM1 among Sudanese population.

## 2. Materials and Methods

### 2.1 Patients

100 patients already diagnosed with AML which was diagnosed using morphology, cytochemistry, immunophenotyping and genetic analysis. EDTA blood or Bone marrow samples were collected from the patients, 34 samples were Bone marrow and 66 samples were peripheral blood sample Clinical and laboratory data, including CBC, French–American–British (FAB) sub class, blast percentage, leukocytes count, Red blood cell count, and hemoglobin (Hb) level were also collected., the patients grouped as 54 were newly diagnosed recruited at the time of diagnosis prior to chemotherapy achievement and 46 were under treatment who attend the RICK as follow up, accordingly, All of them were screened for NPM1 mutations.

### 2.2. Methods

DNA was extracted using manual guanidine technique

#### 2.2.1. Analysis of the NPM1mutations

The extracted DNA were analyzed for detection of NPM1 mutation on chromosome 5q35.1 exon 12 using conventional PCR. 2μl of genomic DNA was amplified in 10μl reaction mixture containing 2μl ready to load mastermix (containing FIRpol DNA polymerase, 5x reaction buffer B, 12.5 Mm MgCl, 1Mm for each deoxyribonucleotidetriphosphate-dNTP) and 1μl of each of the forward and reverse primers. Primers sequences used for NPM1 mutations were mentioned in table No (1). PCR amplification was performed using PCR thermal cycler. Amplification process consisted of initial denaturation at 95C^0^ for 2 mins, 30 cycles of 30 sec at 95C^0^ for denaturation, 58.5C^0^ for 30 sec for annealing. 50 sec at 72C^0^ for extension and 72C^0^ for 5 mins as final extension. Bioinformatic analysis was done using (Mega 7 software programe).

**Table No (1):**
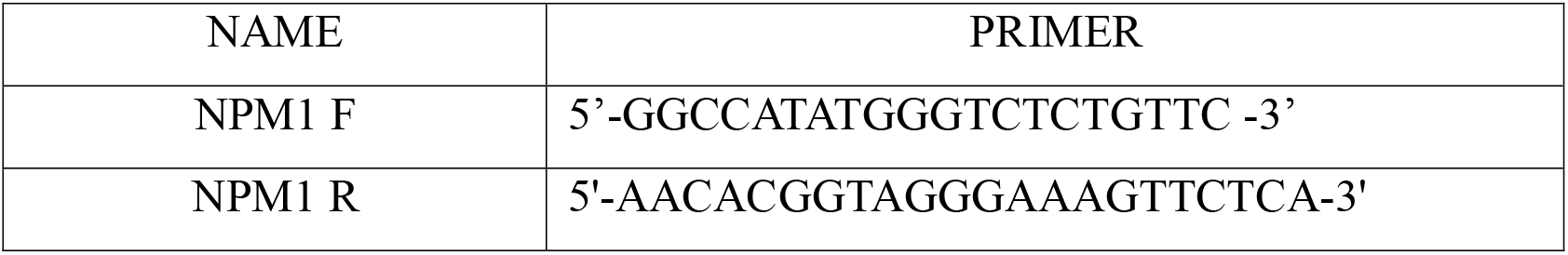
Shows the primers of NPM1 used in the study.

## 3. Results

This study included One hundred patients diagnosed as AML based on morphological, immunophenotype, and molecular genetics with acute myeloblastic leukemia; 54 of them were males and 46 were females; the mean of age is (39 years). (23%) of the patients were FAB type M0 as shown in figure (1). NPM1 mutations detection, sequencing technique was done, then sequencing analysis by software (Mega 7 Software) revealed that there were no NPM1mutations in Sudanese AML patients shown in figure (2).

**Figure (1):**
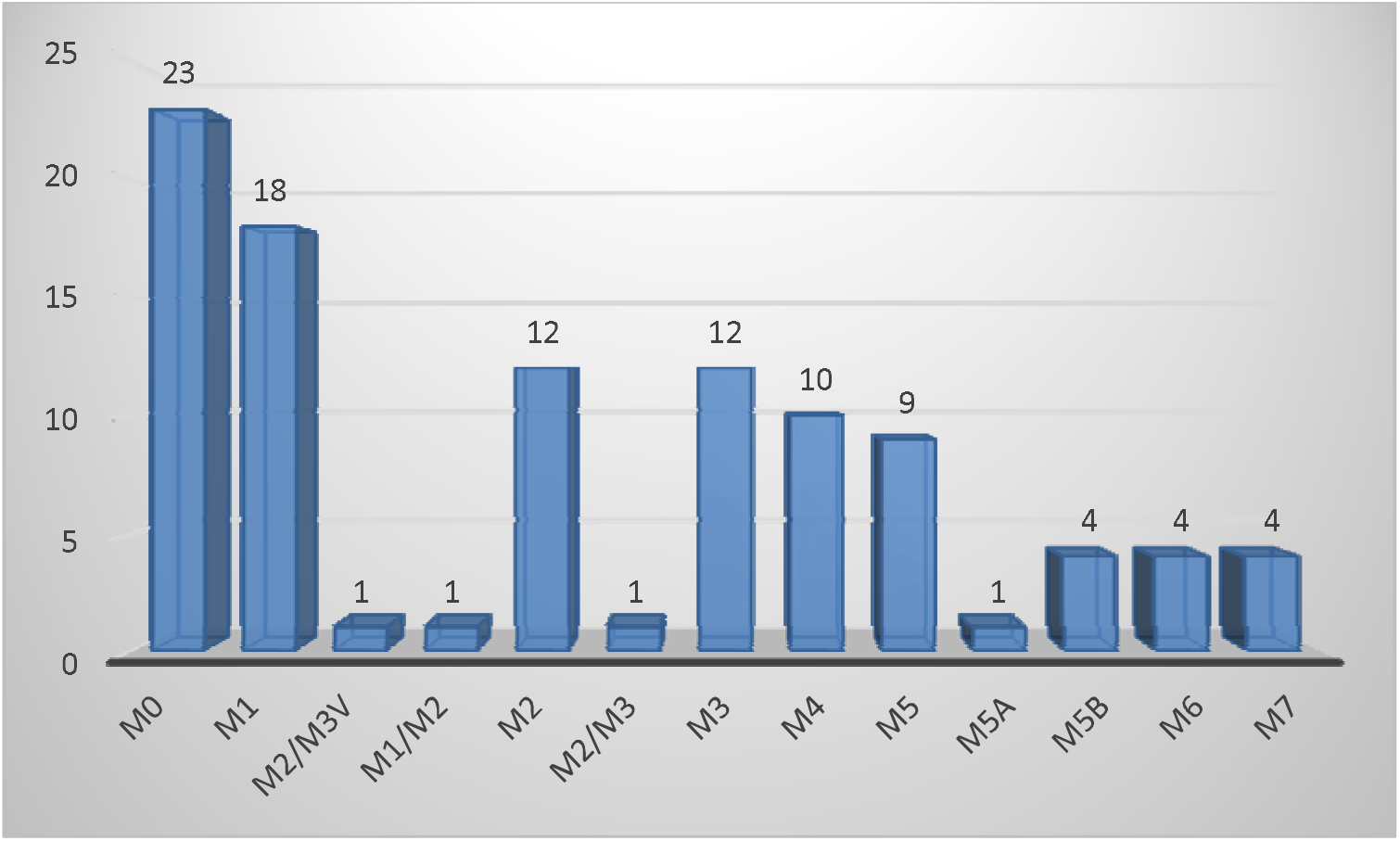
Shows the percentage of AML sub types in the patient of this study.

**Figure (2):**
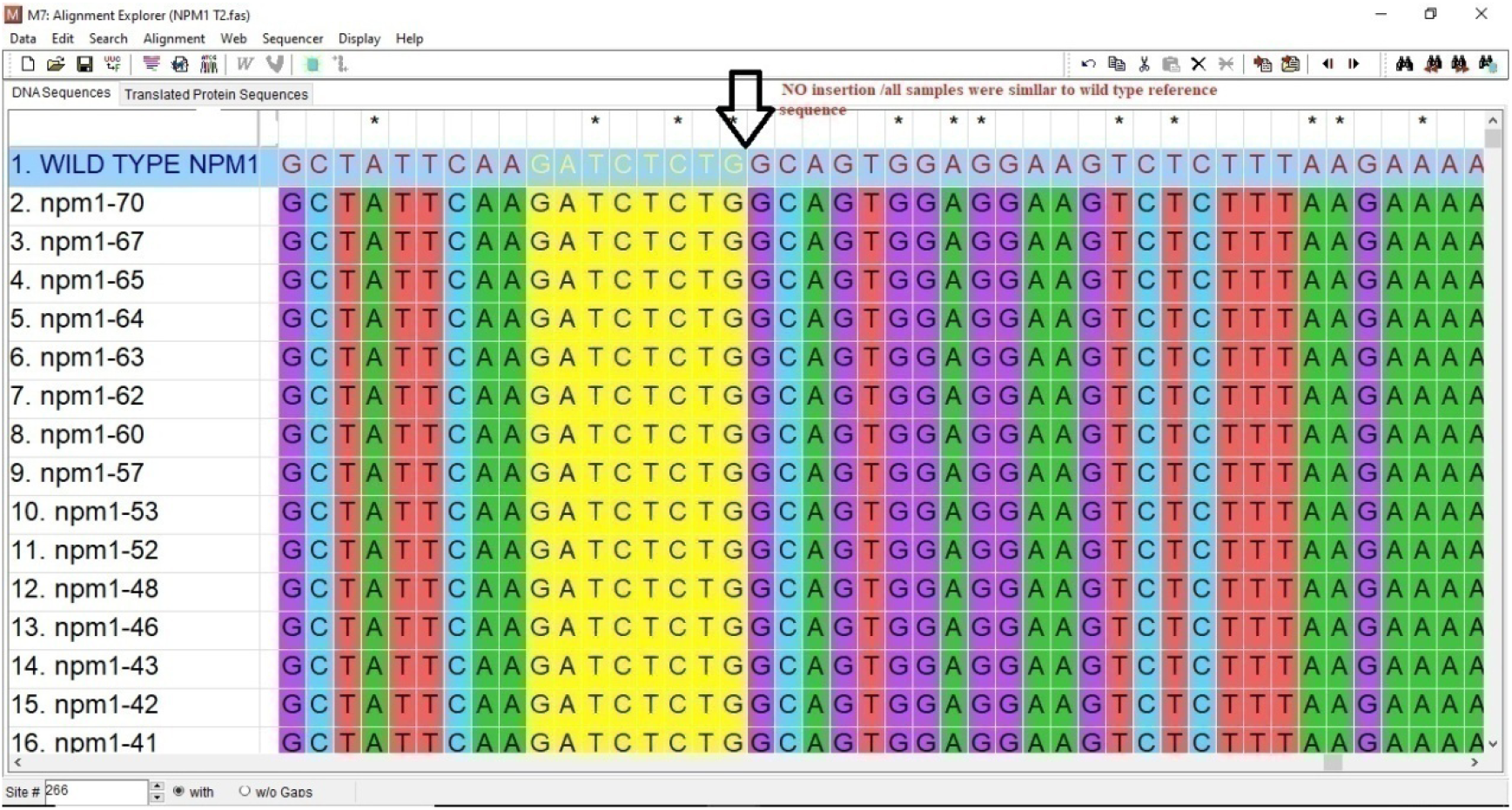
Shows the alignment of NPM1 sample number (20) with wild Types reference.

## 4. Discussion

NPM1 gene characterize as candidate for prognosis and treatment of AML are which revealed that the Sudanese AML patient carry wild type NPM1 which were almost resemble previous study done in Sudan showed minimum prevalence of mutated NPM1 types (3%) (Tariq **et al** 2018 unpublished data) this may due different technique used QRT. PCR comparing to PCR/sequencing which may indicating of methylation effect on the gene (epigenetic effect) rather than nucleotide changes, Therefore when comparing the phenotypes result of Sudanese NPM1 wild types with previous studies from other population revealed were the same figure (Hernández et al, 2013).

One of explanation AML samples used for NPM1 typing missing the karyotype data, may be most of samples were abnormal karyotypes.

Therefore the absence of mutant types of NPM1 among Sudanese population may due to weak role of these mutant types on AML prognosis this low genetic variability due founder effect on this founder effect or genetic drift.

Mutations of NPM1 gene are significantly associated with increasing age and it has been reported that they are less frequently seen in patients under the age of 35 years (Verhaak *et al*, 2005). As more than 50% of the Sudanese patients were 35years or less, this could possibly be the reason behind the absence of the NPM1 mutations in this study. Another possible explanation is that different frequencies have been reported in different ethnic regions, and also globally found significantly lower frequency of NPM1 mutations in Asian populations, for example, 11% in a large study from China (Shen *et al*, 2011).

## 5. Conclusion

NPM1 mutations are part of genetic abnormalities of Acute myeloid leukemia (AML) mainly for diagnosis, treatment but for prognosis they have no clear role in Sudanese patients.

